# Paraoxonase and acylated homoserine lactones in urine from patients with urinary tract infections

**DOI:** 10.1101/641688

**Authors:** John Lafleur, Richard L. Amdur

## Abstract

Paraoxonases are mammalian enzymes that have a number of roles including the inhibition of bacterial virulence and biofilm formation by microorganisms that quorum sense with acylated homoserine lactones. Paraoxonases have previously been reported to inhibit *P. aeruginosa* biofilm formation in mammalian airways and skin. An innate immune role for paraoxonases in urinary tract infection has not previously been reported. We performed western blots for paraoxonase1 in urine from patients with urinary tract infection; we also tested urinary tract infection urine for the presence of acylated homoserine lactones using a cellular reporter system. We report here that paraoxonase1 was not found with our western blot assay in the urine of normal control patients; in those with urinary tract infection, paraoxonase1 was associated with E. *coli* UTI. Acylated homoserine lactones, but not paraoxonases, were found in the bulk urine of those with *P. aeruginosa* urinary tract infection. We hypothesize that paraoxonase may play a similar innate immune role in infected urine as has previously described in skin and airways.

## Introduction

The paraoxonase PON family of mammalian lactonases are an evolutionarily conserved (1–3) innate immune mechanism that limit bacterial virulence and biofilm formation by degrading quorum sensing (QS) acylated homoserine lactones (AHLs) produced by some microorganisms.(4–13) These bacteria include *P. aeruginosa* as well as other environmental opportunists with large genomes and flexible lifestyles that are frequently found to be occult members of infecting biofilms.(14–16) Many of these belong to the group of non-fermenting gram negative bacilli (NFGNB).(17,18) UTI caused by NFGNB have been reported to more commonly infect those with urinary catheters, diabetes mellitus, and previous hospitalizations.(19) In multiple studies over the course of the last decade, PON have been shown to be protective against infection by *P. aeruginosa* biofilm formation in mammalian airways and skin cells.(9,20) Another body surface that is subject to environmental exposure is the urinary system, and PON has previously been reported in urine from healthy subjects.(21)

AHLs are known to be potent promoters of biofilm formation and virulence expression in gram negative pathogens that quorum sense with them, while at the same time causing tissue inflammation and derangement of host immunity.(22–26) Although they have not previously been reported, the presence of AHLs in urine from patients with UTI would support a role for PON in the defense against infection by *P. aeruginosa*.

Uncomplicated UTI in an immunocompetent host is characterized by single species infection, 26 *E. coli* 80% of the time. It may be hypothesized that the limited spectrum of uncomplicated UTI uropathogens is due to competent immunity, including the inhibition by PON of environmental opportunists that QS with AHLs, such as *P. aeruginosa* and many of the NFGNB. To further explore these issues, we set out to assay for AHLs and PON in urine from patients with UTI presenting to the busy emergency department of a large city hospital. We hypothesized that 1) UTI patients will have PONS present, while non-UTI patients will not; 2) among UTI patients, the presence of PONS will be significantly associated with the presence of urinary pathogens

## Materials and methods

### Human subjects enrollment

Study protocol was reviewed/approved by Lifspan IRB

#### Study site

Anderson emergency department of Rhode Island Hospital

#### Inclusion criteria

1. Greater than 18 years of age and able to give informed consent for study participation.
2. 10 or more white cells in urine analysis with symptoms of urinary tract infection.
3. Urine culture sent to the hospital microbiology department (prior to administration of antibiotics).

#### Control subjects

Emergency department patients with minor complaints unrelated to urinary system and without significant metabolic derangement such as fever, hyperglycemia, renal disease (acute or chronic), significant hypertension.

#### Urine samples

once enrolled in study subjects were asked to provide 50-100 ml of clean-catch urine in a sterile cup. This was immediately frozen at −80 for further study.

#### Growth media

Plates and broth were Luria-Bertani (LB).

#### Bacterial culture

The long chain HSL reporter strain *E. coli* JM109 (pSB1142) (carries *P. aeruginosa las*R fused to luxCDABE), and *P. aeruginosa* PAO1 carrying PlasB-luxCDABE were grown in LB broth with shaking at 38 deg. C.

#### Reagents

3-oxo-C12-HSL stock solution 20 mg/ml (Sigma-Aldrich) was diluted 1:50,000 in water. This dilution was arrived at empirically by testing against luminescence in the long chain HSL reporter strain *E. coli* JM109.

#### Western blotting

Urine samples from enrolled research subjects with UTI were stored at −80 deg. C., and thawed for use. 25 microliter samples of unprocessed urine were assayed for PON1 using the Bio-Rad iBlot system as previously described.

#### Antibodies

Primary antibody: polyclonal human PON1 from rabbit (Atlas Antibodies).

#### Secondary antibody

goat anti-rabbit reporter

### Measures

Culture results were recoded into a binary variable (positive/negative). As a sensitivity analysis, we also coded those patients who were positive but with <50k cfu as negative. Positively skewed continuous variables and those with outliers were recoded into ordinal variables.

### Data analysis

Associations between diagnosis and categorical variables were analyzed using chi-square or Fishers Exact Test. Comparison of continuous variables across groups was done using 2-tailed independent groups t-tests or the Kruskal-Wallis test for skewed variables. In order to test the independent association of PON1 with being culture-positive in UTI patients, we used multivariable logistic regression, adjusting for variables that might be confounds. These were defined as having an association with PON1 with p<.10. We also used a multivariable logistic regression model to develop an optimal prediction model for being culture-positive in UTI patients. This was based on the patient variables that were associated with being culture-positive with p<.10, dropping any for which an odds ratio could not be calculated due to low sample size. SAS version 9.4 (Cary, NC) was used for data analysis, with p<.05 considered significant.

## Results and discussion

For the 70 patients in the study, mean age was 60 ± 22, 13 (19%) were black and 44 (63%) were white, and 48 (69%) were female. 11 (16%) were catheterized. Culture was positive in 39/61 cases (64%), while PON1 was positive in 22 cases (36%). There were 61 UTI patients and 9 controls in the sample.

Controls and UTI patients differed significantly on age, with UTI older, on serum creatinine (UTI higher), on highest temperature (UTI higher), and on lowest DBP (UTI lower), and there was a trend-level association for hemoglobin (UTI lower) (Table 1). Patients with UTI were older, and, probably for that reason, have higher average creatinine--due to age-related decline in kidney function (Table 1). Higher temperature in UTI subjects is likely due to some subjects being systemically ill.

**Table 1.** Patient variables by diagnosis (UTI vs control)

| Patient variable | Control (n=9) | UTI (n=61) | p |
| --- | --- | --- | --- |
| Age | 42 ± 16 | 63 ± 22 | .009 |
| Race |  |  | .45 |
| Black | 3 (33%) | 10 (16%) |  |
| White | 5 (56%) | 39 (64%) |  |
| Other | 1 (11%) | 12 (20%) |  |
| Gender female | 4 (44%) | 44 (72%) | .13 |
| Catheterized | 0 | 11 (18%) | .34 |
| WBC | 9.3 ± 4.4 | 11.4 ± 4.4 | .25 |
| Hemoglobin | 13.7 ± 0.9 | 12.5 ± 1.9 | .07 <sup>A</sup> |
| Serum creatinine | 0.8 ± 0.1 | 1.2 ± 0.9 | .042 <sup>A</sup> |
| Highest HR | 85 ± 14 | 93 ± 21 | .29 |
| Highest temp | 97.7 ± 0.6 | 99.0 ± 1.5 | .0001 |
| Lowest systolic bp | 123 ± 16 | 118 ± 20 | .40 |
| Lowest diastolic bp | 78 ± 11 | 68 ± 13 | .044 |

<sup>A</sup> using Kruskal-Wallis test.

PON1 was significantly associated with UTI diagnosis. Of the 61 UTI patients, 22 (36%) were PON1 positive, while none of the controls were PON1 positive (Fisher Exact test p=.049). PON1 was not significantly associated with any demographic or laboratory values (Table 2), but was significantly associated with higher HR (higher in PON1 positive) (Table 2).

**Table 2.** Associations between patient variables and PON1 in patients with UTI

| Patient variable | PON1 Neg (n=39) | PON1 Pos (n=22) | p |
| --- | --- | --- | --- |
| Age | 62 ± 22 | 63 ± 21 | .82 |
| Race |  |  | .60 |
| Black | 5 (13%) | 5 (23%) |  |
| White | 26 (67%) | 13 (59%) |  |
| Other | 8 (21%) | 4 (18%) |  |
| Gender female | 29 (74%) | 15 (68%) | .61 |
| Catheterized | 5 (13%) | 6 (27%) | .18 |
| WBC | 11.2 ± 4.4 | 11.7 ± 4.6 | .68 |
| Hemoglobin | 12.5 ± 2.0 | 12.4 ± 1.7 | .75 |
| Serum creatinine | 1.2 ± 1.0 | 1.2 ± 0.5 | .48 <sup>A</sup> |
| Highest HR | 88 ± 16 | 101 ± 25 | .03 |
| Highest temp | 99.0 ± 1.6 | 98.9 ± 1.3 | .66 |
| Lowest systolic bp | 120 ± 20 | 114 ± 20 | .27 |
| Lowest diastolic bp | 68 ± 12 | 70 ± 13 | .45 |

PON1 was significantly associated with positive culture in UTI patients: PON was positive in 4/22 with negative culture (18%) versus 18/39 with positive culture (46%; p=.03; Table 3). We did a sensitivity analysis coding patients who had culture < 50k cfu as negative (rather than the default coding as positive), and found that the association was still significant (culture negative had 23% PON positive, culture positive had 48% PON positive, p=.04). Thus, in UTI patients, presence of PON in urine was associated with urine culture growing out a urinary pathogen, in contrast to urogenital flora, or no growth. However, in the sample of UTI patients, after adjusting for higher HR, the association between PON1 and culture was no longer significant (an OR for PON1 Pos vs Neg was 3.08 [95% ci 0.84-11.20], p=.09).

**Table 3.** Association of Positive Culture with PON1 and other patient variables, in patients with UTI. N and column-% are shown.

| Culture | Culture Neg<br>(n=22) | Culture Pos<br>(n=39) | p |
| --- | --- | --- | --- |
| PON1 Positive | 4 (18%) | 18 (46%) | .03 |
| Age | 60 ± 24 | 64 ± 20 | .45 |
| Race |  |  | .17 |
| Black | 1 (5%) | 9 (23%) |  |
| White | 16 (73%) | 23 (59%) |  |
| Other | 5 (23%) | 7 (18%) |  |
| Gender female | 16 (73%) | 28 (72%) | .94 |
| Catheterized | 1 (5%) | 10 (26%) | .045 |
| WBC | 9.8 ± 3.5 | 12.2 ± 4.6 | .046 |
| Hemoglobin | 12.7 ± 1.7 | 12.3 ± 2.0 | .44 |
| Serum creatinine | 1.1 ± 0.6 | 1.2 ± 1.0 | .44 |
| Highest HR | 86 ± 16 | 97 ± 22 | .064 |
| Highest temp | 98.8 ± 1.4 | 99.0 ± 1.5 | .61 |
| Lowest systolic bp | 119 ± 19 | 116 ± 21 | .58 |
| Lowest diastolic bp | 71 ± 9 | 67 ± 14 | .23 |

In addition to PON1, other patient variables that were associated with being culture-positive, in UTI patients, included WBC (higher with culture positive), and being catheterized (more frequent for culture positive). Highest HR was marginally associated with culture-positive (Table 3). We created an optimal prediction model for being culture-positive, which included PON1, highest HR, and WBC (being catheterized was dropped because OR could not be calculated for this variable due to small sample size), which had an area under the ROC curve of 0.72 for predicting culture-positive.

Using the equation: risk = −2.43 + 1.007*PON1 + .014*highestHR + .138*WBC, and then probability = exp(risk) / (1 + exp(risk)), and then splitting the probabilities into tertiles, we found that the observed incidence of being culture positive in tertiles 1 through 3, respectively, were 39%, 75%, and 83% (p=.01).

19 out of 38 positive urine cultures grew out E. coli alone (50%). PON1 was positive in 10 of these (53%). When compared to cultures that grew out multiple organisms (including those with ‘urogenital flora’) PON was significantly associated with cultures that grew out E. coli alone, P=0.05 (Table 4). Four NFGNB (10%) were cultured from PON negative urines (all of them *P. aeruginosa*). In addition, three other gram-negative environmental opportunists were cultured from PON negative urines: *Serratia marscesens, Citrobacter freundii*, and *Klebsiella pneumoniae*. These organisms are lactose fermenters, so do not meet criteria for being NFGNB, however, like NFGNB, they are multi-drug resistant environmental opportunists. Additionally these bacteria have all been reported to produce or QS with AHLs.(28–30) Among PON positive urines there were no NFGNB in culture.

**Table 4.** Univariable association of presence of PON in urine with E. coli alone versus cultures growing out multiple different bacteria (including ‘urogenital flora’)

|  | **E. coli alone**<br>(n=21) | **Mult. organisms.**<br>(n=25) |
| --- | --- | --- |
| PON pos. | 10 (53%) | 6 (24%) |
| PON neg. | 9 (47%) | 19 (76%) |

<sup>A</sup> Chi-square 5.3; P=0.05

Using an E. coli luminescent reporter construct, C12 AHL was also assayed for in urine samples. C12 AHLs were only seen in the urines that were culture positive for *P. aeruginosa*. Concentrations of C12 AHL in one sample (patient #9) was about 1.5 micromolar. The other three samples in which *P. aeruginosa* grew out of culture had considerably lower concentrations (see table 5). Biologically relevant concentrations of AHLs for QS are considered to be 1-5 micromolar.(25) One possible interpretation of concentrations of C12 considerably below this in three of four samples suggests that QS and virulence expression in *P. aeruginosa* UTI is not a planktonic phenomenon in the urine but occurring on mucosal surfaces of the bladder/urinary system where surface colonization/biofilm formation can take place, and immune interactions are likely to take place.

**Table 5.** Subjects whose cultures grew out *P. aeruginosa* had urine that contained C12 AHL (except for subject #61 where it was not detectable).

| **Subject #** | **Luminescence (ALU)** | **Micromolar conc.** |
| --- | --- | --- |
| 9 | 167,000 | 1.5 |
| 25 | 22,000 | 0.2 |
| 35 | 8,000 | 0.07 |
| 61 | 0 | 0 |

These preliminary results indicate a positive association between PON level and positive culture in patients with UTI, although HR may be functioning as a confounding factor. Levels of C12 AHL in planktonic cultures necessary to initiate QS-related *las*B expression have previously been reported to be about 1 micromolar(31), a concentration of C12 AHLs that is not uncommonly seen in planktonic cultures of *P. aeruginosa.* By contrast, C12 AHL levels associated with *P. aeruginosa* biofilms in flow cells have been found to be hundreds of times higher.(32) C12 AHL levels in urine from subjects with *P. aeruginosa* UTI have not previously been reported, though detection in urine of non-AHL *P. aeruginosa* mediators of QS associated with pulmonary infection has recently been reported.(33) In the current study results are somewhat equivocal as 3 out of 4 urines were found to contain concentrations of C12 AHLs significantly below the 1 micromolar threshold. There are two possible scenarios that may be imagined in the case of *P. aeruginosa* UTI—that QS occurs mostly on bladder mucosa surfaces where concentrations of microorganisms are likely to be higher, and interactions with mediators of host immunity more intense, or that QS is also occurring in bulk urine. Host bladder epithelial response to UTI include urination, exocytosis of intracellular urinary pathogens, and sloughing of bladder epithelial cells with adherent or intracellular urinary pathogens.(34) Since most planktonic *P. aeruginosa* in UTI will be flushed with urination, possible adaptive advantages conferred by QS among planktonic population of *P. aeruginosa* are that production of virulence factors, such as C12 AHL, may be ramped up in the *P. aeruginosa* population as a whole if QS is occurring in bulk urine. The results of the current study are preliminary, but since 3 of 4 samples showed what appear to be significantly sub-threshold levels of C12 AHL, QS at mucosal surfaces seems more likely. Urine has recently been reported to independently promote *P. aeruginosa* biofilm formation,(35) suggesting that in the absence of the normal QS-mediated mechanisms for biofilm formation, a biofilm may still be formed in *P. aeruginosa* UTI.

We found in the current study that the presence of PON predicts growth of uropathogens as opposed to “urogenital flora” or, “no growth”. The latter two are results of urine culture which are not considered to represent significant infections, but, rather, some source of inflammation resulting in urinary symptoms mimicking acute UTI. One possible explanation for this is that UTI-mediated upregulation of urinary system TLR4/5 may result in the increases expression of PON seen in our PON positive UTI subjects. Urinary TLR signaling has been found to be sensitive to uropathogens,(36,37) resulting in activation of NF-kB and the expression of the pro-inflammatory genes IL-6 and IL-8 with consequent ingress to the bladder mucosa of neutrophils.(38) There is no report in the literature of TLR4/5 mobilization of PON, so at present this remains conjecture. By combining information on PON1 level, higher HR, and WBC, it was possible to obtain an accurate estimate of the probability of having positive cultures.

Our findings include that PON positive subjects had significantly more UTIs caused by *E. coli* alone, rather than multi-species infections, or infections with opportunists such as *P. aeruginosa.* It has previously been reported that the large majority of uncomplicated UTIs in normal hosts are caused by single species—meaning that under normal conditions community richness in UTI is very limited compared to other human microbiomes.(27) Infection associated with impaired immunity is characterized by difficult to eradicate biofilms, polymicrobial infections, and infection with opportunistic organisms that don’t readily infect immune-competent hosts. Viewed in this light, PON positive patients more nearly conform to immune-competent patients with UTIs caused by a single pathogen, while PON negative patients were more likely to have UTIs more characteristic of deranged immune competence. The current study does not make it possible to draw any causal link between the presence of PON in urine and the immune-competent pattern of uncomplicated UTI caused by a single pathogen, mostly *E. coli*. However, the role of PON in innate immunity of the airway and skin,(9,20) and the role that AHL QS plays in many NFGNB environmental opportunists, along with possible occult roles played by fastidious environmental opportunists in establishing complex multi-species infections,(39) suggests that urinary PON may have a protective role.

## Conclusion

The current study reports for the first time AHLs in the urine of subjects with *P. aeruginosa* UTIs. However, the significance of this, and the role that AHLs play in QS among planktonic *P. aeruginosa* remains to be investigated. We found that UTI subjects with PON positive urines were much more likely to have uncomplicated *E. coli* UTI. What mechanism, if any, underlies this finding is at present unclear.

